# A natural variant and an engineered mutation in a GPCR promote DEET resistance in *C. elegans*

**DOI:** 10.1101/198705

**Authors:** Emily J. Dennis, May Dobosiewicz, Xin Jin, Laura B. Duvall, Philip S. Hartman, Cornelia I. Bargmann, Leslie B. Vosshall

## Abstract

DEET (N,N-diethyl-meta-toluamide) is a synthetic chemical, identified by the United States Department of Agriculture in 1946 in a screen for repellents to protect soldiers from mosquito-borne diseases^1,2^. Since its discovery, DEET has become the world’s most widely used arthropod repellent^3^, and is effective against invertebrates separated by millions of years of evolution, including biting flies^4^, honeybees^5^, ticks^6^, and land leeches^4,7^. In insects, DEET acts on the olfactory system^5,8–14^ and requires the olfactory receptor co-receptor orco^9,11–13^, but its specific mechanism of action remains controversial. Here we show that the nematode *Caenorhabditis elegans* is sensitive to DEET, and use this genetically-tractable animal to study its mechanism of action. We found that DEET is not a volatile repellent, but interferes selectively with chemotaxis to a variety of attractant and repellent molecules. DEET increases pause lengths to disrupt chemotaxis to some odours but not others. In a forward genetic screen for DEET-resistant animals, we identified a single G protein-coupled receptor, *str-217*, which is expressed in a single pair of DEET-responsive chemosensory neurons, ADL. Misexpression of *str-217* in another chemosensory neuron conferred strong responses to DEET. Both engineered *str-217* mutants and a wild isolate of *C. elegans* carrying a deletion in *str-217* are DEET-resistant. We found that DEET can interfere with behaviour by inducing an increase in average pause length during locomotion, and show that this increase in pausing requires both *str-217* and ADL neurons. Finally, we demonstrated that ADL neurons are activated by DEET and that optogenetic activation of ADL increased average pause length. This is consistent with the “confusant” hypothesis, in which DEET is not a simple repellent but modulates multiple olfactory pathways to scramble behavioural responses^12,13^. Our results suggest a consistent motif for the effectiveness of DEET across widely divergent taxa: an effect on multiple chemosensory neurons to disrupt the pairing between odorant stimulus and behavioural response.

---

We used standard chemotaxis assays^15–17^ (Fig. 1a) to explore whether and how *C. elegans* nematodes respond to DEET. There are currently three competing hypotheses for the mechanism of DEET: “smell-and-repel” —DEET is detected by olfactory pathways that trigger avoidance^5,10,14,18^, “masking” —DEET selectively blocks olfactory pathways that mediate attraction^8–10^, and “confusant” —DEET modulates multiple olfactory sensory neurons to scramble the perception of an otherwise attractive stimulus^12,13^.

**Figure 1.**
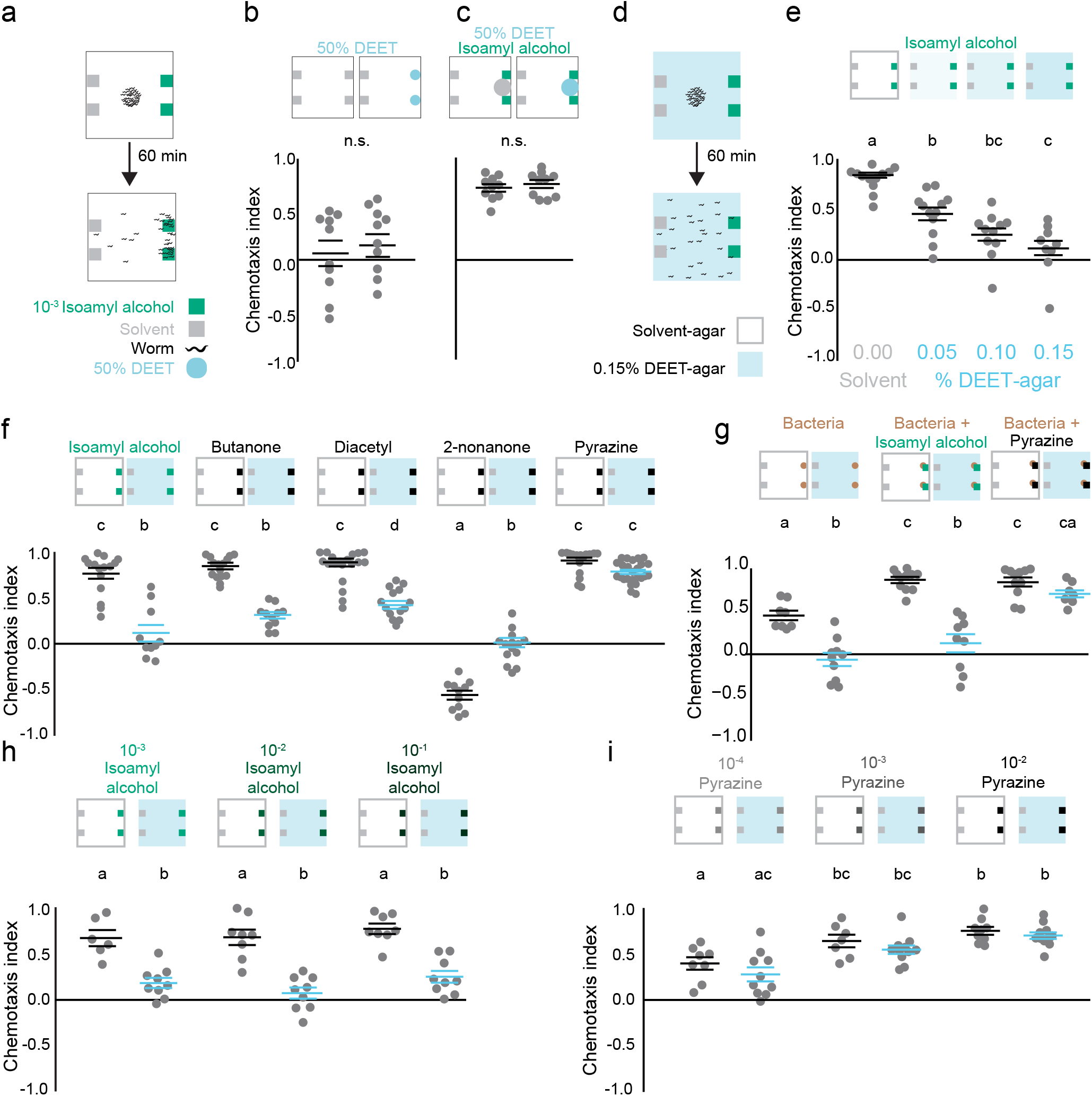
DEET interferes with *C. elegans* chemotaxis. **a**, Schematic of chemotaxis assay. **b-c**, Chemotaxis with point source stimuli of DEET alone (b) or DEET with isoamyl alcohol (c) (N=10). **d**, Schematic of chemotaxis assay on DEET-agar plates. **e**, Chemotaxis to isoamyl alcohol on DEET-agar plates of the indicated concentrations (N=10-13). **f**, Chemotaxis on solvent-agar (grey) or DEET-agar (blue) in response to the indicated odorants (N=11-24). **g**, Chemotaxis to bacterial food source on the indicated plates with indicated odorants (N=7-11). **h-i**, Chemotaxis dose-response curves on the indicated plates to isoamyl alcohol (h) (N=6-9) and pyrazine (i) (N=7-11). In b-c, and e-i, each dot represents a chemotaxis index of a single population assay (50-250 animals). Data labelled with different letters indicate significant differences [mean ± s.e.m , p<0.05, Student’s T-test (b-c) or one-way (e) or two-way (f-i) ANOVA and Tukey’s Post-hoc test].

To test the smell-and-repel hypothesis, we presented DEET as a volatile point source. DEET was not repellent alone (Fig. 1b), similar to previous results in *Drosophila melanogaster* flies^9^ and *Aedes aegypti* mosquitoes^13^, but in contrast to results from *Culex quinquefasciatus*^19^. To address the possibility that DEET could be masking responses to attractive odorants^8,9^ or directly inhibiting their volatility^10^, we presented DEET alongside the attractant isoamyl alcohol, both as point sources, and found that DEET had no effect on attraction (Fig. 1c). In considering alternate ways to present DEET, we mixed low doses of DEET uniformly into the chemotaxis agar and presented isoamyl alcohol as a point source (Fig. 1d). In this configuration, DEET-agar reduced chemotaxis to isoamyl alcohol in a dose-dependent manner (Fig. 1e). To determine if DEET has a general effect on chemotaxis, we tested three additional attractants, butanone, diacetyl, and pyrazine, as well as the volatile repellent 2-nonanone. DEET decreased attraction to butanone and avoidance of 2-nonanone, indicating that it can affect responses to both positive and negative chemosensory stimuli (Fig. 1f). DEET also affected chemotaxis to diacetyl, but not pyrazine, which is notable because both odorants require the same primary sensory neurons, AWA (Fig. 1f). A similar effect is seen in *D. melanogaster*, where DEET can affect responses to some odours and not others, even in a single chemosensory neuron^20^.

In *D. melanogaster* flies and *Ae. aegypti* mosquitoes, DEET inhibits attraction to complex odour blends of food and host odours^21,22^. When we provided the food odour of bacterial suspensions of OP50 *E. coli* to *C. elegans*, we found that DEET eliminated chemotaxis to this complex odour blend (Fig. 1g). Remarkably, supplementing bacterial odour with pyrazine, but not isoamyl alcohol, restored chemotaxis (Fig. 1g). To exclude the possibility that pyrazine is able to overcome the effect of DEET due to a higher effective concentration, we carried out dose-response experiments with isoamyl alcohol (Fig. 1h) and pyrazine (Fig. 1i) and found that at all concentrations tested, DEET interfered with isoamyl alcohol chemotaxis but not pyrazine chemotaxis. From these data we conclude that DEET chemosensory interference is odour-selective, and can affect both attractive and repellent stimuli.

Identifying genes required for DEET-sensation has been of interest for some time. A forward genetic screen in *D. melanogaster* yielded an X-linked DEET-insensitive mutant^23^, and a population genetics approach in mosquitoes identified a dominant genetic basis for DEET-insensitivity^24^, but neither study identified the underlying genes. Reverse genetic experiments in *D. melanogaster* and three mosquito species identified orco, the co-receptor for insect odorant receptor genes, as a molecular target of DEET^9,11–14^. However, this chemosensory gene family is not found outside of insects^25,26^, raising the question of what pathways are required for DEET-sensitivity in non-insect invertebrates. To gain insights into the mechanisms of DEET repellency in *C. elegans*, we carried out a forward genetic screen for mutants capable of chemotaxing toward isoamyl alcohol on DEET-agar plates (Fig. 2a). We obtained 5 DEET-resistant animals, three of which produced offspring that consistently chemotaxed toward isoamyl alcohol on DEET-agar plates (Fig. 2b, and data not shown). We identified candidate causal mutations in two strains using whole genome sequencing^27^, and focused on *LBV003*, which maps to *str-217*, a predicted G protein-coupled receptor (Fig. 2c and d). In the course of mapping *str-217*, we discovered that a divergent strain of *C. elegans* isolated in Hawaii, CB4856 (Hawaiian), is naturally resistant to DEET (Fig. 2e). This Hawaiian strain contains a 138-base pair deletion in *str-217 (str-217^HW^)* that affects exons 2 and 3 and an intervening intron, leading to a mutant strain with a predicted frame shift and early stop codon (Fig. 2c and d and Supplemental Data File 1). This is not a unique phenomenon. Using published data from the *C. elegans* Natural Diversity Resource (CeNDR)^28^, we discovered that 119 of 247 sequenced strains contain predicted changes in the STR-217 protein when compared to N2 wild-type (Supplemental Data File 1). One of these mutant strains shares the *str-217^HW^* deletion and 13 have a missense mutation that introduces a stop codon after the fourth amino acid. The remaining 105 strains have one or more of 30 high-confidence non-synonymous substitutions. We further explored DEET resistance in the Hawaiian deletion by testing three near-isogenic lines with a single, homozygous genomic segment of Hawaiian chromosome V introgressed into a wild-type (Bristol N2) background^29^ (Fig. 2e). Only the ewIR74 line contains *str-217^mv^* and, like the parent Hawaiian strain, is DEET-resistant (Fig. 2e). To provide further confirmation that *str-217* is required for DEET sensitivity in these strains, we generated an additional genetic tool, a rescue/ reporter plasmid that expresses both wild-type *str-217* and green fluorescent protein (GFP) under control of a predicted *str-217* promoter (Fig. 2f). The *LBV003* mutant strain (Fig. 2g) and the Hawaiian introgressed strain *ewIR74* (Fig. 2h) both showed chemotaxis on DEET-agar. The *str-217* rescue/reporter construct rendered both DEET-resistant mutants fully sensitive to DEET, in that neither chemotaxed to isoamyl alcohol on DEET-agar (Fig. 2g and h).

**Figure 2.**
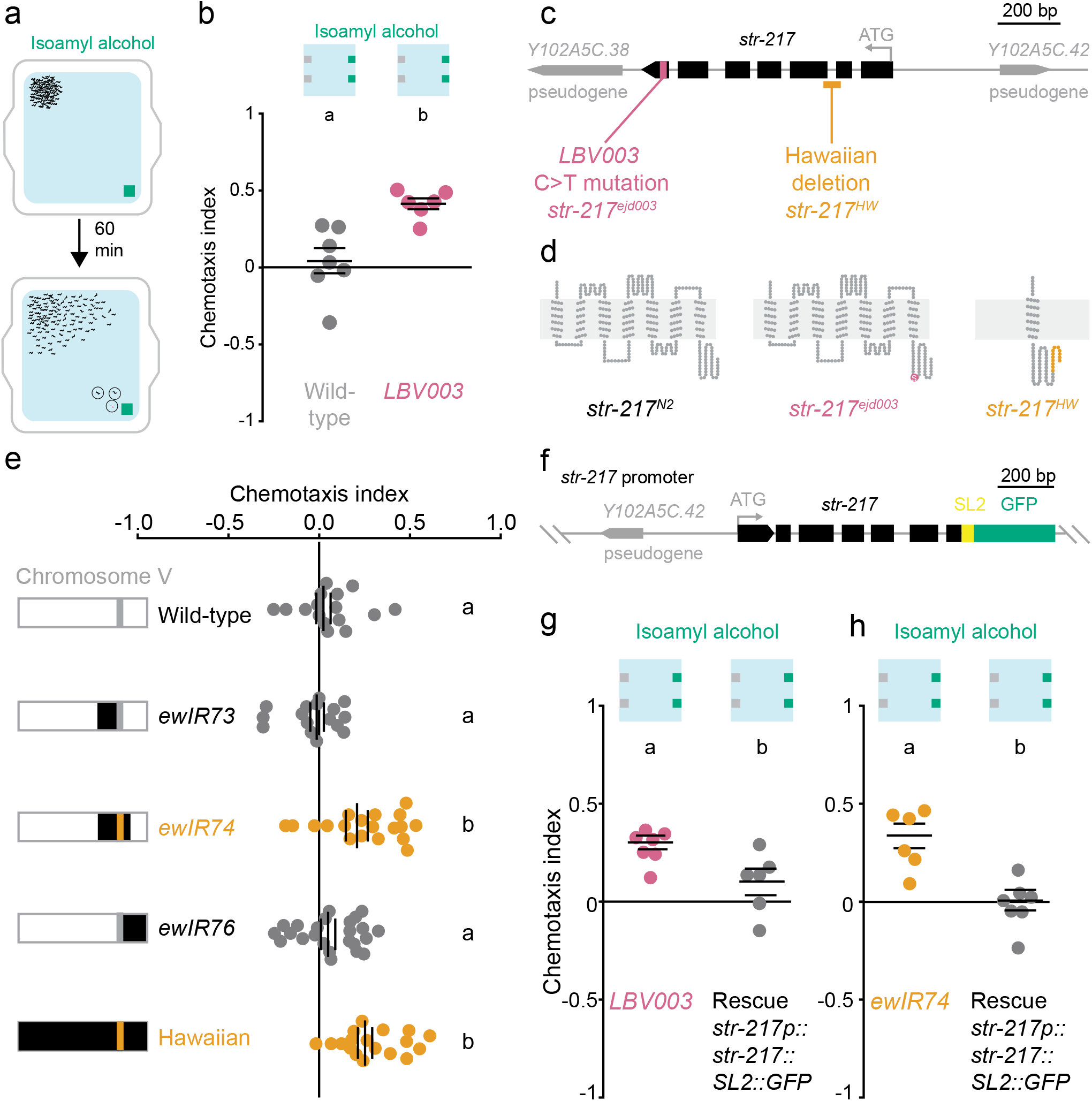
*str-217* mutants are DEET-resistant. **a**, Schematic of forward genetic screen with hypothetical DEET-re-sistant mutants circled. **b**, Chemotaxis of the indicated strains (N=7,6). **c**, Annotated *str-217* genomic locus. **d**, Snake plots of predicted STR-217 proteins in the indicated strains. **e**, Left: schematic of chromosome V in the indicated strains. Right: chemotaxis of the indicated strain (N=16-24). **f**, Schematic of *str-217* rescue/reporter construct. **g-h**, Chemotaxis indices of the indicated strains (N=6-9). In b, e, and g-h each dot represents a chemotaxis index of a single population assay (50-250 animals). Data labelled with different letters indicate significant differences (mean ± s.e.m.; p<0.05 ANOVA and Tukey’s Post-hoc test in e, and two-sided Student’s t-test in b and g-h).

We next turned to the neuronal mechanism by which DEET disrupts chemotaxis in *C. elegans*. In insects, DEET interacts directly with chemosensory neurons and the odorant receptors that they express^9,11–14^. Isoamyl alcohol is primarily sensed by AWC chemosensory neurons^30^. To investigate if DEET modulates primary sensory detection of isoamyl alcohol, we used *in vivo* calcium imaging to monitor AWC odour responses in the presence and absence of DEET. AWC responded to the addition of DEET with a rapid increase in calcium that decreased to baseline over the course of 11 min of chronic DEET stimulation (Fig. 3 a). In the presence of DEET, AWC responses to isoamyl alcohol decreased in magnitude, but there were no significant differences in AWC activity between wild-type and *str-217^−/-^*, a predicted null *str-217* mutant produced by CRISPR-Cas9 genome-editing (Fig. 3a-c). This suggests that AWC sensory neurons are not the primary functional target of DEET, even though they are affected by it. The polymodal nociceptive neurons ASH also responded to DEET (Fig. 3d and e), but animals lacking ASH are fully DEET-sensitive (Fig. 3f), suggesting that these sensory neurons are also not the primary target of DEET.

**Figure 3.**
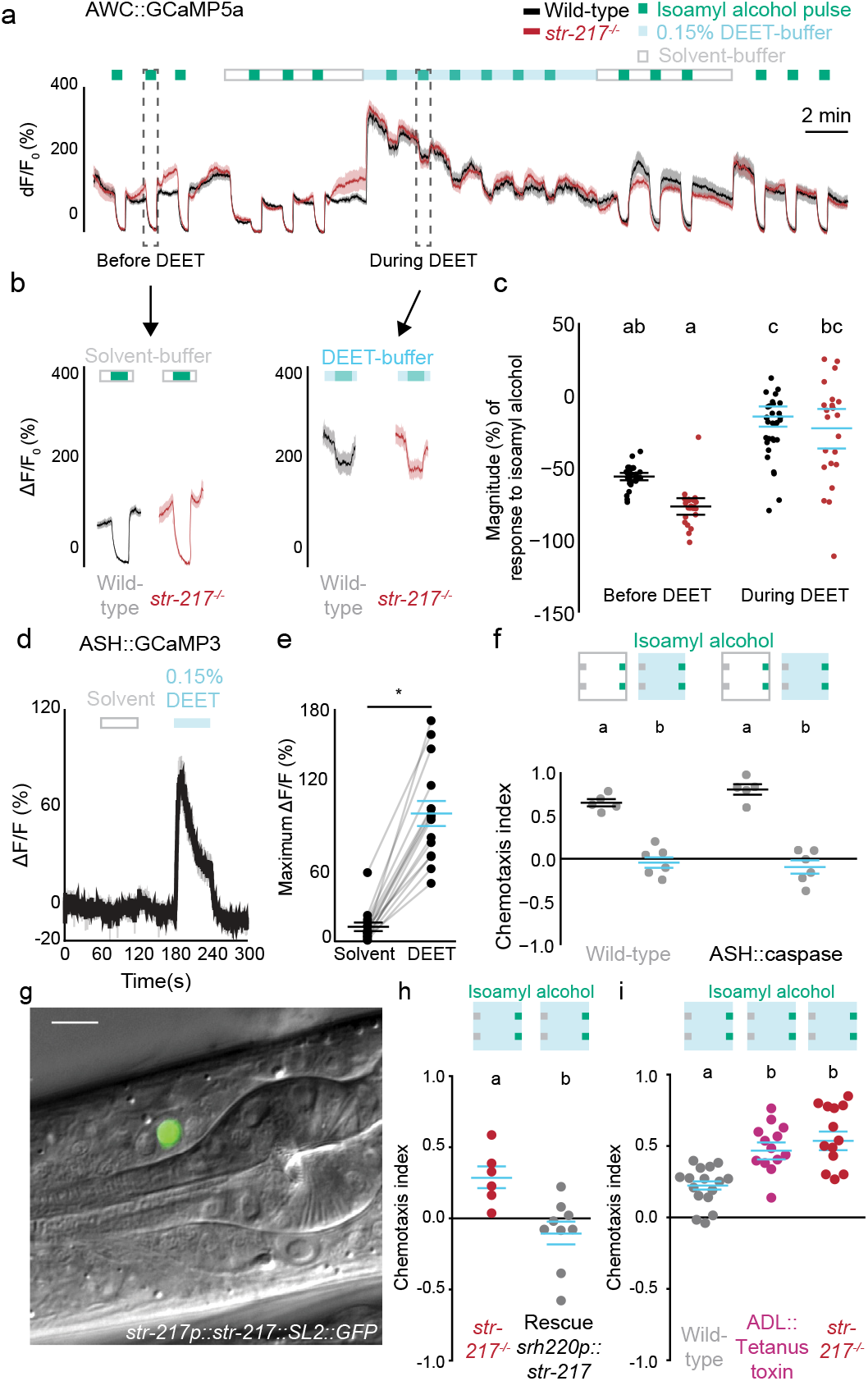
Several *C. elegans* neurons respond to DEET, but ADL neurons are required for DEET-sensitivity. **a**, Average traces of GCaMP activity in AWC^ON^ cells of the indicated genotype over a 36 min experiment. **b**, A subset of traces from (a) re-plotted separately **c**, Response magnitudes of the data in b (N=23 wild-type, 31 *str-217^−/-^*). **d**, GCaMP activity in ASH neurons in response to the indicated stimuli. **e**, Quantification of data in d. **f**, Chemotaxis of the indicated strains to isoamyl alcohol. **f**, GFP expression in a single ADL neuron from *str-217* rescue/reporter construct (scale bar: 10 μm). **h-i**, Chemotaxis of the indicated strains to isoamyl alcohol on DEET-agar. In a-b and d, each trace represents the mean (dark line) and s.e.m. (lighter area) GCaMP response of all animals of each genotype tested. In c and e, each dot represents response of a single animal. In f and h-i, each dot represents a chemotaxis index of a single population assay (50-250 animals) [mean ± s.e.m , p<0.05, two-way (b, e) or one-way (h) ANOVA and Tukey’s Post-hoc test, and a paired Student’s t-test in d and g].

To identify such neurons, we determined where *str-217* is expressed by examining the *str-217* rescue/reporter strain, and found GFP expression in a single pair of chemosensory neurons, called ADL (Fig. 3g). This was unexpected because ADL is not required for chemotaxis to isoamyl alcohol^31^, suggesting an indirect role for ADL in DEET chemosensory interference. To investigate if *str-217* is only required in ADL, we expressed *str-217* in ADL under control of the *srh-220* promoter in *str-217^−/-^* worms. ADL-specific rescue of *str-217* rendered this mutant sensitive to DEET (Fig. 3h). To ask if ADL neuronal function is required for DEET-sensitivity, we inhibited chemical synaptic transmission by expressing the tetanus toxin light chain in ADL^32,33^. These animals showed the same level of DEET-resistance as *str-217* mutants (Fig. 3i), demonstrating that ADL neurons are required for DEET-sensitivity.

Since both *str-217* and ADL function are required for DEET-sensitivity, we used calcium imaging to see if ADL responds to DEET, and if this requires *str-217* (Fig. 4a). Both wild-type and *str-217^−/-^* mutants carrying a rescue plasmid, but not *str-217^−/-^* mutants, showed calcium responses to DEET, albeit with some variability in response (Fig. 4b-d). To exclude the possibility that DEET affects ADL indirectly by interacting with other sensory neurons that subsequently activate ADL, we measured calcium responses of ADL in genetic backgrounds that disrupt chemical synaptic transmission between neurons. ADL neurons showed normal responses to DEET in both *unc-13* and *unc-31* mutant animals, which are deficient in synaptic vesicle fusion^34^ or dense-core vesicle fusion^35^, respectively (Fig. 4f-h). In control experiments, we showed that a known ADL agonist, the pheromone C9^32^, activated ADL in both wild-type, *str-217^−/-^* mutant, and rescued animals (Fig. 4j-l), suggesting that the *str-217^−-^* mutation has a selective effect on ADL responses to DEET.

**Figure 4.**
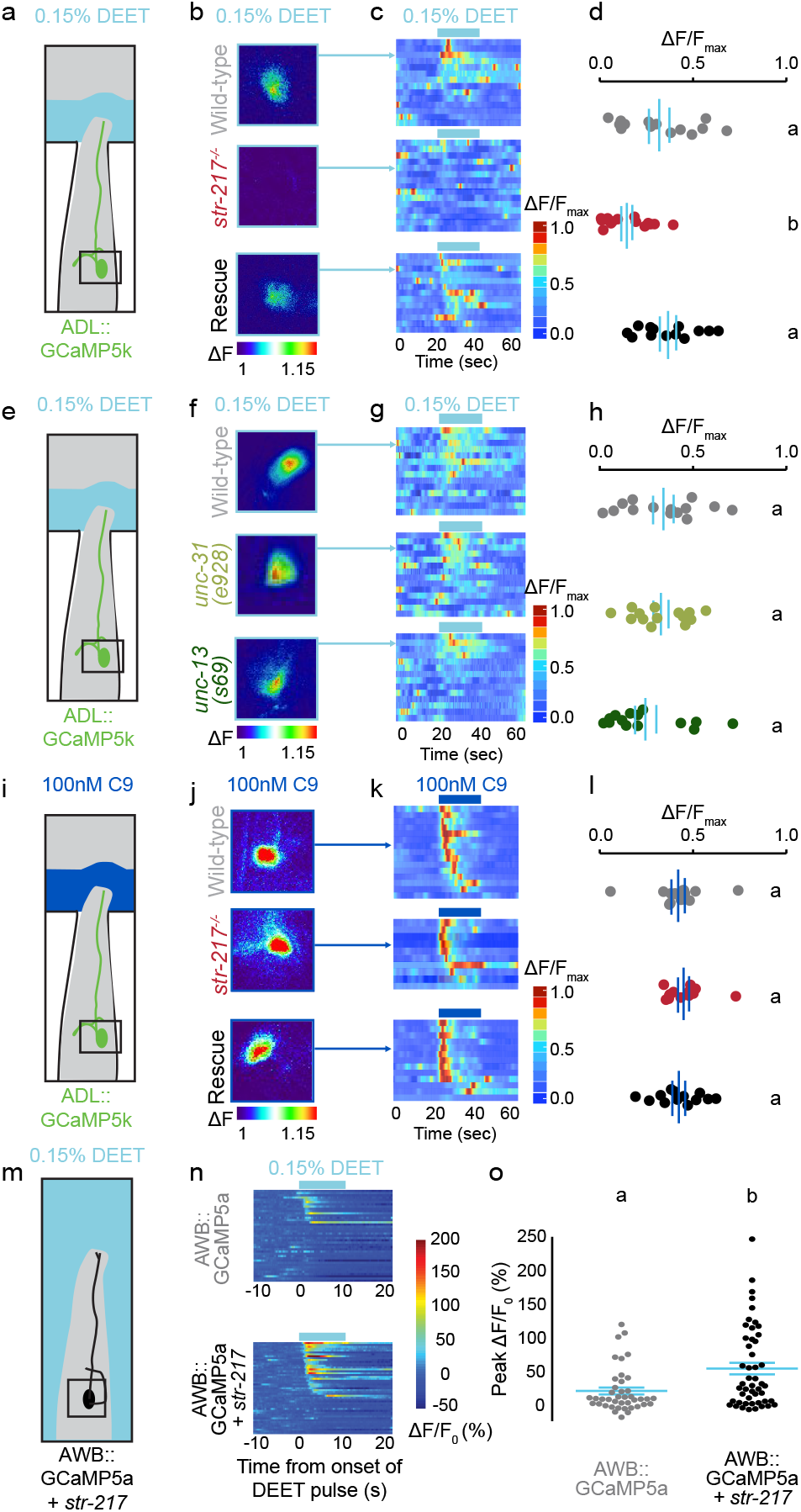
*str-217* is necessary for ADL responses to DEET and can confer DEET sensitivity to AWB. **a, e, i**, and **m**, Schematics of microfluidic calcium imaging assays. **b, f**, and **j**, Pseudocolored images of ADL responding to 0.15% DEET (b, f) or 100 nM C9 (j) in animals of the indicated genotype (fold increase in mean fluorescence 20 s after first stimulus pulse / mean of 20 s before the stimulus pulse). Arrows point to corresponding animal in following panels. **c, g** and **k**, Heat maps of GCaMP activity in response to DEET (c, g) or C9 (k). Each row represents ADL activity in one animal. **d, h**, and **l**, Mean normalized responses of the data in **c, g**, and **k** during DEET (d, h) or C9 (l) 30 second pulse. **n**, Heat maps of calcium imaging data of AWB neurons in response to 0.15% DEET, each row represents imaging from one animal cropped to show the 10 s before, during, and after the DEET pulse. o, Quantification of data in n. In d, h, l, and o each dot represents a single neuron in a single animal. Data labelled with different letters indicate significant differences [mean ± s.e.m. p<0.05, one-way ANOVA and Tukey’s Post-hoc test (d, h, l), or Student’s t-test (o)].The rescue construct used in this figure is the same as the one used in Fig. 3g and h.

Since *str-217* is necessary for ADL to respond to DEET, we asked if it *str-217* is sufficient to confer DEET responses when heterologously expressed. When we expressed *str-217* in HEK293T cells we failed to see activation by DEET (Supplemental Data File). We therefore misexpressed *str-217* in another pair of *C. elegans* chemosensory neurons, AWB, and found that *str-217* significantly increased the DEET-sensitivity of this cell (Fig. 4m-o). These gain-of-function data are consistent with the hypothesis that *str-217* itself encodes a DEET receptor, or cooperates with other proteins *in vivo* that respond to DEET.

We next explored how ADL activity can interfere with chemotaxis. Population chemotaxis assays report the location of the animal at the end of the experiment, but do not reveal the details of navigation strategy. To investigate which aspects of chemotaxis are affected by DEET, we tracked the position and posture of individual animals on DEET-agar or solvent-agar plates (Fig. 5a-c). Wild-type, but not *str-217^−/-^* mutants (Fig. 5d), showed a dramatic increase in average pause length on DEET-agar. Although *str-217^−/-^* mutants are resistant to DEET compared to wild type, their chemotaxis is still reduced to some degree by DEET (Fig. 3h). This is evident in the end-point position of *str-217^−/-^* animals, many of which never make it to the odour source (Fig. 5e). Chemotaxis indices did not increase for wild-type animals when we prolonged the assays (Fig. 5f). This indicates that there are additional effects of DEET on chemotaxis in addition to increased pause duration.

**Figure 5.**
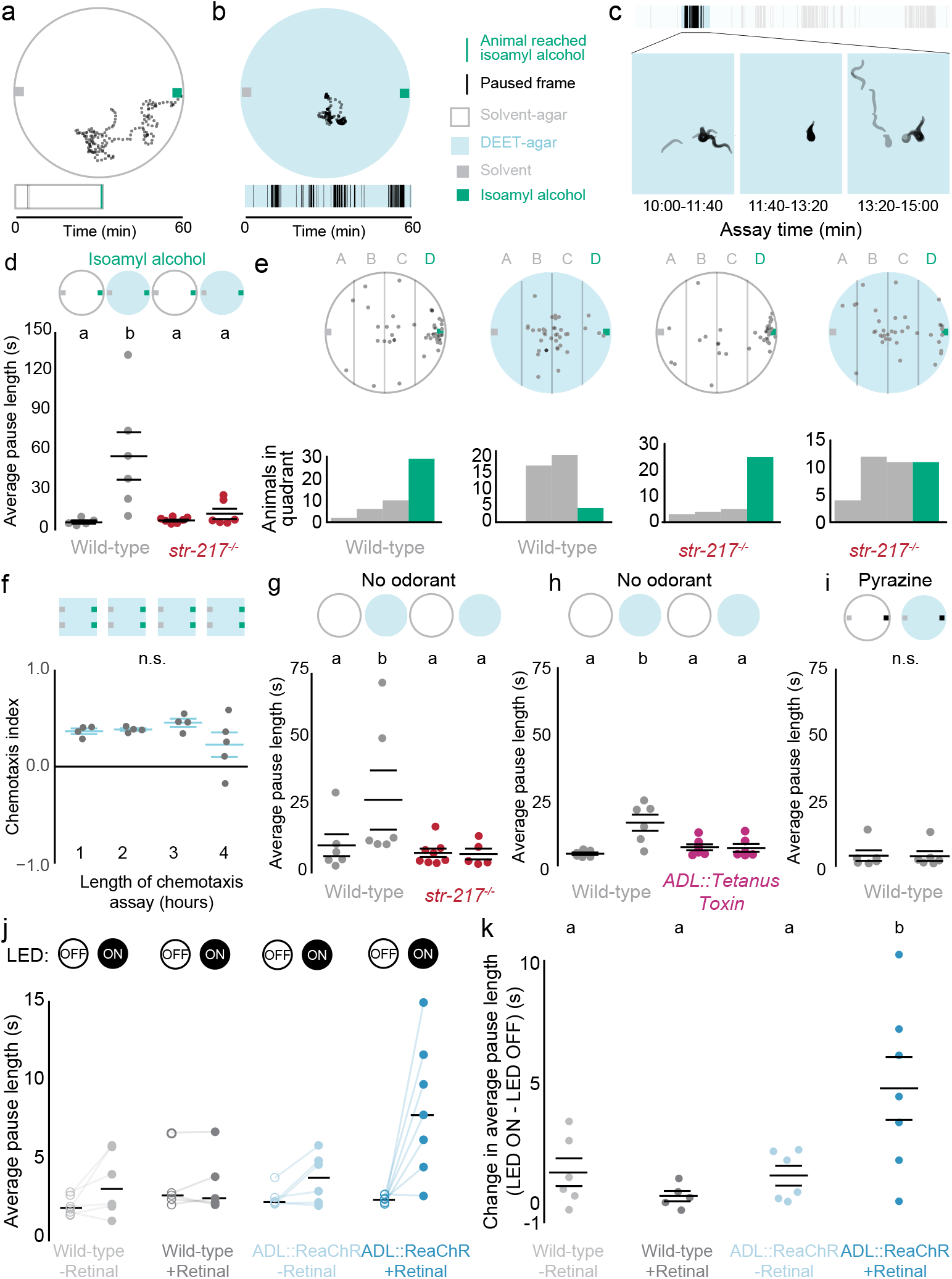
DEET increases average pause length by activating *str-217* and ADL. **a-b**, Top: example trajectories of single wild-type animals chemotaxing to isoamyl alcohol on solvent-agar (a) or DEET-agar (b). Each dot depicts the x, y position of a single animal once every 10 s. Bottom: raster plots of paused frames for the animal above. **c**, Example pauses from animal in b. Images were extracted and converted to silhouettes once every 18 frames (6 s) and superimposed. **d**, Average pause length with the indicated stimuli and genotypes (N=6-7 plates, 4-15 animals per plate). **e**, Top: Location of animals from experiments in d at the end of the 60 minute assay. Bottom: Histogram of animal location from e. **f**, Wild-type DEET chemotaxis indices at indicated assay time. **g-i**, Average pause length of the indicated stimuli and genotypes (N=6-7 plates, 4-15 animals per plate). j, Average pause length of the indicated genotype and LED status, with lines connecting each experimental pair, i, Difference in average pause length for each experimental pair in j (N=6 experiments, 4-15 animals per experiment). In d and f-k each dot represents a single experimental plate of 50-250 animals (f) or 4-15 animals (d, g-k). Data labelled with different letters indicate significant differences [mean ± s.e.m. p<0.05, one-way (f) or two-way ANOVA (d,g-k) and Tukey’s Post-hoc test].

To determine if the increase in average pause length occurs only in the context of chemotaxis to isoamylalcohol, or as a consequence of DEET alone, we tracked wild-type, *str-217^−/-^* mutant (Fig. 5g), and *ADL::Tetanus toxin* (Fig. 5h) animals on DEET-agar and solvent-agar plates with no additional odorants. Only wild-type animals had a higher average pause length on DEET-agar (Fig. 5g-h). Consistent with our prior observation that chemotaxis to pyrazine was unaffected by DEET, wild-type animals showed no increase in average pause length when chemotaxing to pyrazine on DEET-agar (Fig. 5i). This suggests that pyrazine chemotaxis can overcome the effect of DEET on average pause length.

To test if ADL activity alone is sufficient to increase average pause length, we carried out an optogenetic experiment by expressing the light-sensitive ion channel ReaChR^36^ in ADL in wild-type animals, and tracking locomotor behaviour on chemotaxis plates. We observed an increase in average pause length when ADL was activated (Fig. 5j and k). From these data, we conclude that ADL mediates the increase in average pause length seen on DEET-agar, and speculate that the increase in long pauses is one mechanism by which DEET interferes with chemotaxis.

In this study, we add the nematode *C. elegans* to known DEET-sensitive animals and uncover a new neuronal mechanism for a DEET-induced behaviour. We identify *str-217* as a molecular target that is required for DEET-sensitivity in an engineered mutant and in a wild-isolate of *C. elegans*. This work opens up *C. elegans* as a system to test new repellents *in vivo* and also for discovery of additional genes and neurons that respond to DEET. The molecular mechanism by which the *str-217* mutation renders ADL DEET-insensitive and worms DEET-resistant remains to be understood. *str-217* encodes a G protein-coupled receptor with no known ligand and that is evolutionarily unrelated to DEET-sen-sitive odorant receptor proteins previously described in insects. Our results are consistent with the hypothesis that *str-217* is a DEET receptor, or that it interacts with additional proteins to make ADL DEET-sensitive. Inter-estingly, pyrazine chemotaxis is unaffected by DEET in any of our assays, consistent with our model that DEET is not a simple repellent, but a modulator of behaviour that interferes with chemotaxis to some but not all odorants.

Although *str-217* has no clear orthologues outside of nematodes, the *str-217*-dependent mechanism of action of DEET in nematodes we have discovered is reminiscent of the “confusant” hypothesis in insects. In insects, DEET alters responses of individual olfactory sensory neurons to attractive odorants^9,12^, thereby interfering with behavioural attraction. In *C. elegans*, DEET inhibits attraction to some odours by activating neurons that induce competing behaviours like pausing. We speculate that its promiscuity in interacting with multiple molecules and chemosensory neurons across vast evolutionary scales is the key to the broad effectiveness of DEET.

## Acknowledgements

We thank Michael Crickmore, Kevin Lee, Aakanksha Singhvi, Nilay Yapici, and members of the Vosshall Lab for discussion and comments on the manuscript. Shai Shaham advised and Wendy Wang assisted with chemical mutagenesis. Heeun Jang provided guidance on chemotaxis behaviour and imaging. Alejandro Lopez-Cruz and Elias Sheer provided advice on tracking behaviour. Sagi Levy shared the *Pstr-2::GCaMP5a* strain and Elias Sheer shared the *Psrh-220::ReaChR* plasmid. We thank Anh Nguyen for her contributions to the early analysis of DEET-resistant mutants in the Hartman laboratory. Some *C. elegans* strains used in this paper were obtained from the CGC, which is funded by NIH Office of Research Infrastructure Programs (P40 OD010440). This work was supported by an NIH grant to E.J.D. (F31 DC014222). L.B.V. is an investigator of the Howard Hughes Medical Institute.

## Author Contributions

E.J.D. and L.B.V. developed the concept and designed the experiments. E.J.D performed all experiments and analysis unless noted. M.D. performed AWC and ASH calcium-imaging experiments in Figure 3 and the AWB calcium-imaging experiments in Figure 4. X.J. carried out imaging experiments in Figure 4 along with E. J.D., and performed cell identification in Figure 3. L.B.D. performed HEK293T heterologous expression experiments. C.I.B. provided guidance and experimental design advice and interpreted data. P.S.H. made the original observation that DEET interferes with chemotaxis in *C. elegans*, and initiated genetic screens for DEET-resistant mutants in his laboratory. E.J.D. and L.B.V. together interpreted the results, designed the figures, and wrote the paper with input from the other authors. The authors declare no competing financial interests.

## Methods

### Nematode culture and strains

*C. elegans* strains were maintained at room temperature (22-24°C) on nematode growth medium (NGM) plates (51.3 mM NaCl, 1.7% agar, 0.25% peptone, 1 mM CaCl_2_, 12.9 μM cholesterol, 1mM MgSO_4_, 25mM KPO_4_, pH 6) seeded with *Escherichia coli* (OP50 strain) bacteria as a food source^37,38^. Bristol N2 was used as the wild-type strain. The CB4856 (Hawaiian) strain, harbouring WBVar02076179 *(str-217^HW^)* (http://www.wormbase.org/db/get?name=WBVar02076179;class=variation) and Hawaiian recombinant inbred strains for chromosome V were previously generated^29^. Generation of extra-chromosomal array transgenes was carried out using standard procedures^39^, and included the transgene injected at 50 ng/mL, the fluorescent co-injection marker *Pelt-2::GFP* at 5 ng/ml (with the exception of LBV004 and LBV009, which did not include a co-injection marker), and an empty vector for a total DNA concentration of 100 ng/ml. CRIS-PR-Cas9-mediated mutagenesis of *str-217* was performed as described, using *rol-6* as a co-CRISPR marker^40^. The resulting *str-217* mutant strain [LBV004 *str-217(ejd001)]* results in a predicted frame-shift in the first exon [indel: insertion (AAAAAAA), deletion (CTGCTCCA), final sequence GCGTCGAAAAAAAATTTCAG; insertion is underlined]. The *str-217* rescue construct *(Pstr-217::str-217::SL2::GFP)* used a 1112 nucleotide length fragment 56 nucleotides upstream 5’ of the translation start of *str-217*.

### Microscopy and image analysis

L2-adult stage hermaphrodites were mounted on 1% agarose pads with 10 mM sodium azide (CID 6331859, Sigma-Aldrich, catalogue #S2002) in M9 solution (22 mM KH_2_PO_4_, 42mM Na_2_HPO_4_ 85.6 mM NaCl, 1μM MgSO_4_, pH 6). Images were acquired with an Axio Observer Z1 LSM 780 with Apotome a 63X objective (Zeiss), and were processed using ImageJ.

### Chemotaxis assays

Chemotaxis was tested as described^17^, on square plates containing 10 mL of chemotaxis agar (1.6% agar in chemotaxis buffer: 5 mM phosphate buffer pH 6.0, 1 mM CaCl_2_, 1 mM MgSO_4_)^41^. Additions of either ethanol (solvent-agar) or 50% DEET (CID: 4284, Sigma-Aldrich, catalogue #D100951) in ethanol (DEET-agar) were added after agar cooled to <44°C and just before pouring. A total volume of 300 μL ethanol or DEET in ethanol was added to each 100 mL of agar mixture for all experiments except Figure 1b and c,Figure 5j and k. Plates were poured on the day of each experiment, and dried with lids off for 4 hours prior to the start of the assay. 1 mL 1 M sodium azide was added to two spots on either side of the plate just before beginning the experiment to immobilize animals that reached the odorant or ethanol sources. Three days prior to all chemotaxis experiments, 4-6 L4 animals were transferred onto NGM plates seeded with *E. coli* (OP50 strain). The offspring of these 4-6 animals were then washed off of the plates and washed twice with S-Basal buffer (1 mM NaCl, 5.74 mM K_2_HPO_4_, 7.35 mM KH_2_PO_4_, 5 μg/mL cholesterol at pH 6-6.2)^37^ to remove younger animals, and once with chemotaxis buffer. Immediately before the start of the experiment, two 1 mL drops of odorant diluted in ethanol, or ethanol solvent control, were spotted on each side of the plate on top of the sodium azide spots. 100-300 animals were then placed into the centre of the plate in a small bubble of liquid. The excess liquid surrounding the animals was then removed using a Kim-wipe. 0dorants diluted in ethanol were used in this study at these concentrations unless otherwise noted: 1:1000 isoamyl alcohol (CID: 31260, Sigma-Aldrich, catalogue #W205702), 1:1000 butanone (CID: 6569, Sigma-Aldrich, catalogue #360473), 10 mg/μL pyrazine (CID: 9261, Sigma-Aldrich, catalogue #W401501), 1:10 2-nonanone (CID: 13187, Sigma-Aldrich, catalogue #W2787513). For bacterial chemotaxis assays, 20 μL of either LB media, or OP50 bacterial suspension grown overnight and diluted in LB media to 1.0 OD at 600nm was applied instead of or in addition to odorants. Assays were carried out for 60-90 min at room temperature (22-24°C) between 1pm – 8pm EST with the exception of Figure 5f, which were quantified after either 55-65 minutes (1 hour), 115-125 minutes (2 hours), 175-185 minutes (3 hours), or 235-245 minutes (4 hours). Plates were scored as soon as possible, either immediately or, if a large number of plates was being scored on the same day, plates were moved to 4°C to immobilize animals until they could be scored. The assay was quantified by counting animals that had left the origin in the centre of the plate, moving to either side of the plate (#Odorant, #Control) or just above or below the origin (#Other), and calculating a chemotaxis index as [#Odorant – #Control] / [#Odorant + #Control + #Other]. A trial was discarded if fewer than 50 animals or more than 250 animals contributed to the chemotaxis index and participated in the assay.

### Mutant screen

About 100 wild-type (Bristol N2) L4 animals were mutagenized in M9 solution with 50 mM ethyl methanesulfonate (CID: 6113, Sigma-Aldrich, catalogue #M0880) for 4 hours with rotation at room temperature. Mutagenized animals were picked to separate 9 cm NGM agar plates seeded with *E. coli* (OP50 strain) and cultivated at 20°C. ~5,000 F2 animals were screened for DEET resistance on 20.3 cm casserole dishes (ASIN B000LNS4NQ, model number 81932OBL11). Five animals across three assays were more than ~2 cm closer to the odour source than the rest of the animals on the plate and were defined as DEET-resistant. This phenotype was heritable in three strains, and each strain was backcrossed to OS1917 for 4 generations. Whole-genome sequencing^27^ was used to map the mutations to regions containing transversions presumably introduced by the EMS, parental alleles of the N2 strain used for mutagenesis, and missing alleles of the wild-type strain OS1917 used for backcrossing^42,43^. LBV003 mapped to a 5 Mb region on chromosome V, which was further mapped to *str-217*. LBV002 mapped to a 6.8 Mb region on chromosome V, which was further narrowed down to a likely candidate gene, *nstp-3(ejd002)*. In LBV002, *nstp-3(ejd002)* contains a T>G transversion of the 141^st^ nucleotide in the CDS, which is predicted to produce a Phe48Val substitution in this predicted sugar:proton symporter. We were unable to map the DEET-resistant mutation(s) in LBV001.

### *str-217* heterologous expression in mammalian tissue culture cells

HEK-293T cells were maintained using standard protocols in a Thermo Scientific FORMA Series II water-jacketed CO_2_ incubator. Cells were transiently transfected with 1 μg each of pME18s plasmid expressing *GCaMP6s, G_q_15*, and *str-217* using Lipofectamine 2000 (CID: 100984821, Invitrogen, catalogue #1168019). Control cells excluded *str-217*, but were transfected with the other two plasmids. Transfected cells were seeded into 384 well plates at a density of 2 x 10^6^ cells/ml, and incubated overnight in FluoroBrite DMEM media (ThermoFisher Scientific) supplemented with foetal bovine serum (Invitrogen, catalogue #10082139) at 37°C and 5% CO_2_. Cells were imaged in reading buffer [Hanks’s Balanced Salt Solution (GIBCO) + 20 mM HEPES (Sigma-Aldrich)] using GFP-channel fluorescence of a Hamamatsu FDSS-6000 kinetic plate reader at The Rockefeller University High-Throughput Screening Resource Centre. DEET was prepared at 3X final concentration in reading buffer in a 384-well plate (Greiner Bio-one) from a 46% (2 M) stock solution in DMSO (Sigma-Aldrich). Plates were imaged every 1 s for 5 min. 10 μL of DEET solution in reading buffer or vehicle (reading buffer + DMSO) was added to each well containing cells in 20 μL of media after 30 s of baseline fluorescence recording. The final concentration of vehicle DMSO was matched to the DEET additions, with a maximum DMSO concentration of 7.8%. Fluorescence was normalized to baseline, and responses were calculated as max ratio (maximum fluorescence level/baseline fluorescence level) (Supplemental Data File 1).

### ADL calcium imaging

Calcium imaging and data analysis were performed as described^44^, using single young adult hermaphrodites immobilized in a custom-fabricated 3 x 3 x 3 mm polydimethylsiloxane (PDMS) imaging chip. *GCaMP5k* was expressed in ADL neurons under control of the *sre-1* promoter^32^ and was crossed into *str-217^−/-^* and the *str-217^−/-^* rescue strain. Imaging of *unc-13* and *unc-31* mutant strains was performed by crossing *ADL::GCaMP5k* expressing animals to the *unc-* strains and selecting for fluorescent, uncoordinated animals. Animals were acclimated to the imaging room overnight on *E.coli* (OP50 strain) seeded plates. All stimuli were prepared the day of each experiment, and were diluted in ethanol to 1000X the desired concentration before being further diluted 1:1000 in S-Basal buffer. Young adult animals were paralyzed briefly in (-)-tetramisole hydrochloride (CID: 27944, Sigma-Aldrich, catalogue #L9756) at 1 mM for 2-5 min before transfer into the chip to paralyze body wall muscles to keep animals stationary during imaging. All animals were pre-exposed to light (470+/-40 nm) for 100 s before recording to attenuate the light response of ADL^45^. Experiments consisted of the following stimulation protocol: 20 s S-Basal buffer, followed by 3 repetitions of 20 s DEET (0.15% DEET and 0.15% ethanol in S-Basal) and then 20 s S-basal buffer. ADL responses desensitize rapidly^31^, so only the first of the three DEET pulses was analysed.

All GCaMP signals were recorded with Metamorph Software (Molecular Devices) and an iXon3 DU-897 EMCCD camera (Andor) at 10 frames/s using a 40x objective on an upright Zeiss Axioskop 2 microscope. Custom ImageJ scripts^17^ were used to track cells and quantify fluorescence. In Figure 4b, f, and j, all frames in 20 s before the DEET pulse were averaged and divided by the average of the frames during the 20 s DEET or C9 pulse to calculate ΔF. In Figure 4c, g, and k, traces were bleach-corrected using a custom MATLAB script and then the 5% of frames with the lowest values were averaged to create F_0_ ΔF/F_0_ was calculated by (F – F_0_)/F_0_ and then divided by the maximum value to obtain ΔF/F_max_^46^. The average value during the stimulus was calculated for each animal and plotted as a single dot in Figure 4d, h, and l. The heatmap traces in Figure 4c, g, and k were also smoothed by 5 frames, such that each data point *n* is the running average of *n-2, n-1, n, n+1, and n+2*.

### AWC, ASH, and AWB calcium imaging

Calcium imaging of freely moving worms and subsequent data analysis were performed as described^46^, using a 3 mm^2^ microfluidic PDMS device with two arenas that enabled simultaneous imaging of two genotypes with approximately 10 animals each. For AWC imaging, we used an integrated line (CX17256) expressing *GCaMP5a* in AWC^ON^ neurons undei control of the *str-2* promoter crossed into *str-217^−^* animals. Adult hermaphrodites were first paralyzed for 80-100 min in 1 mM (-)-tetramisole hydrochloride and then transferred to the arenas in S-Basal buffer. The stimulus protocol was as follows: In S-Basal, three pulses of 60 s in buffer and 30 s isoamyl alcohol, followed by 120 s in buffer. Next, the animals were switched to S-Basal with 0.15% ethanol (solvent buffer) and three pulses of 60 s in buffer and 30 s in isoamyl alcohol in solvent buffer followed by 120 s in solvent buffer before a switch to S-Basal with 0.15% ethanol and 0.15% DEET (DEET buffer). In DEET buffer, animals were given 6 pulses of 60 s in DEET buffer and then 30 s in isoamyl alcohol in DEET buffer, followed by 120 s in DEET buffer before switching to solvent buffer. In solvent buffer, the animals received three pulses of 60 s in buffer and 30 s in isoamyl alcohol in solvent buffer followed by 120 s in solvent buffer before a switch to S-Basal. In S-Basal, the animals received three pulses of 60 s in buffer and 30 s isoamyl alcohol, followed by 60 s in buffer.

Each experiment was repeated 3-4 times over 2-3 days and pooled by strain for analysis (wild-type: 31 animals, 4 experiments, 3 days; *str-217^−/-^*: 23 animals, 3 experiments, 2 days). Images were acquired at 10 frames/s at 5X magnification (Hamamatsu Orca Flash 4 sCMOS), with 10 ms pulsed illumination every 100 ms (Sola, Lumencor; 470/40 nm excitation). Fluorescence levels were analysed using a custom ImageJ script that integrates and background-subtracts fluorescence levels of the AWC neuron cell body (6×6 pixel region of interest). Traces were normalized by subtracting and then dividing by the baseline fluorescence, defined as the average fluorescence of the last 2 s of the first three isoamyl alcohol pulses. The traces in Figure 3a-b were also smoothed by 5 frames, such that each data point *n* is the running average of *n-2, n-1, n, n+1, and n+2*. The response magnitudes in Figure 3c were calculated by taking the mean of the last 2 s of an isoamyl alcohol pulse, subtracting the mean of the 2 s before the isoamyl alcohol pulse (F0), and dividing by this F0. The response magnitudes were calculated for the 5^th^ (0.15% ethanol in S-Basal buffer) and 8^th^ (0.15% DEET and 0.15% ethanol in S-Basal buffer isoamyl alcohol pulses.

ASH calcium imaging was performed similarly with the following exceptions. For ASH imaging, we used a strain (CX10979) expressing *GCaMP3* in ASH neurons under control of the *sra-6* promoter. The stimulus protocol used was as follows: 60 s in S-Basal, 60 s in 0.15% ethanol in S-Basal buffer, 60 s in S-Basal, 60 s in 0.15% DEET in S-Basal, and finally 60 s in S-Basal buffer. Each experiment was repeated over 2 days and pooled for analysis (wild-type: 15 animals in 2 experiments on 2 different days).

AWB calcium imaging was performed similarly to AWC imaging with the following exceptions. For AWB imaging, we used a control strain (CX17428) expressing *GCaMP5a* in AWB neurons under the *str-1* promoter and a test strain (CX17660) expressing *GCaMP5a* under the *str-1* promoter as well as expressing *str-217* in AWB neurons under the *str-1* promoter. Adult hermaphrodites were first similarly paralyzed in 1 mM (-)-tetramisole hydrochloride for 80-100 minutes, but the first 65-75 minutes was in S-Basal buffer and the last 15 minutes was 1 mM (-)-tetramisole hydrochloride in ethanol-buffer. The stimulus protocol used was as follows: 60 s in 0.15% ethanol in S-Basal buffer, 10 s in 0.15% DEET and 0.15% ethanol in S-Basal buffer, and 60 s in 0.15% ethanol in S-Basal buffer. A 4x4 pixel region of interest was used during tracking of the neurons. Baseline fluorescence was defined as the median fluorescence of the 10 s preceding the DEET pulse. Response and peak magnitudes were calculated using traces smoothed by 5 frames and identifying the maximum value within the DEET pulse. Five sets of experiments were conducted over 3 days for a total of 41 wild-type animals and 49 animals expressing *str-217*.

In Figure 4n, traces were bleach corrected using a custom MATLAB script and then the 5% of frames with the lowest values were averaged to create F_0_ ΔF/F_0_ (%) was calculated by (F – F_0_)/F_0_^46^. The heatmap traces in Figure 4n were also smoothed by 5 frames, such that each data point *n* is the running average of *n-2, n-1, n, n+1, and n+2*. Peak ΔF/F_0_ (%) in Figure 4o reflects the maximum value of ΔF/F_0_ (%) during the DEET pulse.

### Chemotaxis tracking and analysis

8-20 adult hermaphrodites were first transferred to an empty NGM plate and then 4-15 were transferred to an assay plate to minimize bacterial transfer. Animals were then placed in the centre on either a 0.15% DEET-agar or solvent-agar plate, and their movement was recorded for 60 min at 3 frames/s with 6.6 MP PL-B781F CMOS camera (Pix-eLINK) and Streampix software. Assays were carried out at room temperature, between 12-8pm EST, and lit from below. Worm trajectories were extracted by a custom Matlab (MathWorks) script^17^, and discontinuous tracks were then manually linked. Tracks were discarded if the animal moved less than two body lengths from its origin over the course of the 60 min trial. If an animal came within 1 cm of the isoamyl alcohol stimulus, the track was truncated to remove information from animals immobilized at the odour source because of the addition of sodium azide.

### ADL optogenetic stimulation

L4 animals expressing an *Psrh-220::ReaChR*^36^ array or array-negative animals from the same plate were raised overnight in the dark on an NGM plate freshly seeded with 100 μL of 10X concentrated *E. coli* (OP50 strain) with or without 50 μM all-trans retinal (CID: 720648, Sigma-Aldrich, catalogue #R2500), which is required for ReaChR-induced activity. The next day, adult hermaphrodites were first transferred to an empty NGM plate and then 4-15 animals were transferred to the 10 cm circular assay plate to minimize bacterial transfer. Videos were recorded for 26 min at 3 frames/s with a 1.3 MP PL-A741 camera (PixeLINK) and Streampix software. Blue light pulses were delivered with an LED (455 nm, 45 μW/mm^2^, Mightex) controlled with a custom Matlab script^17,47^. Animals were exposed to normal light for 120 s, before exposure to 6 repetitions of blue light (10 Hz strobing) for 120 s, and 120 s of recovery (LED OFF). Worm trajectories were extracted by a custom Matlab script^47^. Pausing events were extracted, and all pauses ≥3 frames (1 s) were used for further analysis. Pauses were classified as “ON” if any frame included light illumination. A pause that began just before illumination began, but remained paused while the illumination occurred, was considered an ON pause, as well as any pauses that began in the during light illumination considered ON. All other pauses were classified as “OFF” pauses. In the analysis in Figure 5j, we took an average pause length for all ON pauses and all OFF pauses for each animal and pooled all of the animals on each plate. To control for any baseline differences between animals and experiment-to-experiment variation, we examined the increase in average pause length in Figure 5k.

### Phylogenetic Analysis of str-217 in wild isolates

Data were obtained from CeNDR36^28^ and plotted in R. Only predicted deletions in exons or missense changes of high confidence were included (Supplemental Data File 1).

### Statistical Analysis

R v3.3.2 was used for all statistical analysis. Inclusion and exclusion criteria were pre-established for all experiments, and plate positions were pseu-do-randomized in behaviour experiments. Additionally, qqPlots were evaluated before performing ANOVAs. For the analysis of optogenetic experiments, a Levene’s Test identified heteroskedasticity in these data that was addressed with a boxcox translation. AWC imaging data were similarly boxcox translated and transformed to adjust for the rightward skew. All data necessary to re-create these plots are available in Supplemental Data File 1. Data, scripts to analyse these data, and all statistical analyses are available at GitHub: http://github.com/VosshallLab/Dennis-Emily_2017

### Strains

Detailed genotypes of all C. *elegans* strains and their sources are in Supplemental Data File 1.

